# Spectral features of heart rate variability in Williams syndrome during sleep

**DOI:** 10.1101/2025.11.01.685999

**Authors:** Bence Schneider, Ferenc Gombos, Ilona Kovács, Róbert Bódizs

## Abstract

**Aims:** The study aims to reveal the spectral alterations of heart rate variability (HRV) in Williams syndrome (WS) during sleep, using a two-slope, broken power-law model, taking into account the multi-fractal properties of the RR-interval spectra, including effects of aging and associations with sleep structure indicators.

**Methods & Results:** Extracting ECG recordings from a polysomnography database of 20 subjects with WS and 20 age and sex matched typically developing (TD) controls, RR-interval time series were constructed, then the fractal and oscillatory power-spectral densities computed using the Irregular-Resampling Auto-Spectral Analysis (IRASA) method. The fractal component was parametrized with a piecewise-linear function, that allowed for a custom breaking point in the power spectral density (PSD), and separate slope and intercept values in the lower and higher frequency domains. The frequency and prominence of the dominant peak was extracted from the LF (0.04–0.15 Hz) and HF (0.15–0.4 Hz) bands.

Strong WS vs TD group differences were found in the frequency of the breaking point, high domain slope, intercept and HF peak prominence. Analysis of the LF peak frequency revealed age-dependent decrease in the TD, but not in the WS group, while generally decreased values in WS independent of age, potentially reflected accelerated aging, characteristic of the syndrome. As fractal parameters were correlated, a main component was identified using principal component analysis that described the typical alterations in the fractal spectrum in WS, and was correlated with sleep structure indices as well.

**Conclusions:** The broken power-law model proved to be successful in characterizing the fractal component of HRV spectra, furthermore it captured alterations in cardiac regulation in WS. A main spectral feature of WS was identified in the fractal component, being associated with sleep quality indicators, a possible biomarker of the degree of general autonomic deregulation inherent to the syndrome.

## Introduction

Williams syndrome (WS, also known as Williams-Beuren syndrome) is a multisystem congenital genetic disorder caused by the microdeletion of the chromosome region 7q11.23 that can be described by mild intellectual disability, developmental delay, distinctive facies, altered cognitive abilities that include increased short-term memory and verbal skills, in contrast, deficiencies in visuospatial cognition. Individuals with WS exhibit some specific personality traits like excessive empathy, social disinhibition, and are often comorbid with ADHD and generalized anxiety disorder. [Morris CA, 1999]. Atypical development in WS can be manifested as early onset of puberty, premature graying of the hair, high frequency hearing loss, cataracts, etc., all indicating accelerated aging.

More than half WS patients are affected by sleep disorders, specifically by difficulties in sleep initiation and maintenance, decreased total sleep time and sleep efficiency, while intra-sleep wakefulness and daytime sleepiness are more frequent. The acceleration of age-related sleep deterioration was also observed in several sleep quality indices in WS compared to typically developing subjects. [Bódizs et al., 2014]

The prevalence of cardiovascular diseases is high (80%) in WS, being the largest cause of mortality, with aortic stenosis forming the most typical condition due to the reduced elastic properties of vessel walls as a consequence of elastin insufficiency inherent to the syndrome. [Collins et al, 2018] [Lin et al., 2022]

Even in the absence of severe pathologies, alterations in cardiac regulations can be present, e.g. different arrhythmias were identified without being associated with structural cardiac disease [Deitch et al., 2023]

The coexistence of dysregulations in sleep [Bódizs et al. 2012, 2014, Gombos et al., 2010,], cardiac activity [Collins, 2013] and affective functioning [Järvinen et al., 2012] suggest a general involvement of the autonomic nervous system (ANS) in the symptoms of the syndrome. This is further evidenced by altered heart rate reactivity and electrodermal activity in response to affective social stimuli [Järvinen et al., 2012], no arousal level synchronization with others, indicated by a lack of pupil size contagion [Kleberg et al., 2023], as well as diminished circadian modulation in the heart rate and a general reduction in heart rate variability (HRV) indices, observed in 24-hour ECG recordings [Levin et al., 2023]. Moreover, recent evidence suggests a fragmentation of heart rate of WS subjects on a short time scale, involving erratic behaviour characterized by relatively frequent beat-to-beat accelerations and related emergence of inflection points. Authors suggest the potential relevance of nonautonomic cardiovascular modulators in the emergence of this picture [Cathey et al., 2024].

The HRV index had been considered to reflect changes in the ANS, with the low frequency (LF) 0.04–0.15 Hz component corresponding to sympathetic and the high frequency (HF) 0.15–0.4 Hz component to vagal activation. Besides cardiovascular diseases, changes in HRV have been identified in conditions known to associate with WS, among which ADHD and anxiety disorders [ChucDuc et al., 2013] [Cheng et al., 2022] are of potential interest.

In the context of accelerated aging in WS, the HRV parameters might also provide relevant information, as it was proposed that HRV is not only associated with chronological age, but potentially a biomarker of processes that contribute to aging, and thus it is able to reveal trajectories of unhealthy aging. [Olivieri et al., 2024] Despite the large overlap between the specific sensitivity of HRV indices and WS physiopathology, aspects of the syndrome are underexplored in the HRV analysis framework.

Apart from autonomous control, the heart rhythm is modulated by many factors ranging from the molecular scale to the organ level [Monfredi & Lakatta, 2019], as such the RR-interval time-series comprises a complex, fractal component, which is manifested as a power-law in the power spectral density (PSD) of the time-series [Kobayashi & Musha, 1982]. The exponent of the power-law, or the so-called spectral slope characterizes the persistence and long-term memory of the signal, it was found to carry meaningful information about electrophysiological signals, such as local field potentials, electrocorticograms, electroencephalograms, magnetoencephalograms [Donoghue et al. 2020, Colombo et al., 2019].

The aim of our study is to identify spectral alterations in HRV due to Williams syndrome in sleep, using a parametric model that takes into account the underlying multifractal nature of RR-interval time series, that has been identified as a property of healthy heart rate dynamics [Bigger et al., 1996][Ivanov et al., 1999]. Furthermore, to explore the association between sleep structure and the extracted spectral parameters.

## Methods

Bipolar ECG derivations were analysed from an ambulatory polysomnography database [Bódizs et al., 2012] containing whole-night home sleep recordings (recording lengths range: 6.66–12.19 hours, mean: 10.36 hours) of 20 subjects with WS (13 females, age range: 6–29 years, mean: 19.6 years) and an age and sex matched control group of 20 subjects (14 females, age range: 6–29, mean: 20.3 years). All participants had given their written consent to participate in the data recording; the procedure had been approved by the Ethical Committee of the Budapest University of Technology and Economics, in line with the Declaration of Helsinki. Recordings were performed by using a 32 channel SD-LTM Hardware together with the BRAIN QUICK SystemPLUS EVOLUTION software (Micromed, Italy). Signals were high-pass filtered at 0.33 Hz and low-pass filtered at 1500 Hz by a 40 dB/decade anti-aliasing hardware input filter. Data were collected with 22 bit resolution and with an analogue to digital conversion rate of 4096 Hz/channel (synchronous). A further 40 dB/decade anti-aliasing digital filter was applied by digital signal processing which low-pass filtered the data at 124 Hz. After this the digitized and filtered EEG was subsequently undersampled at 1024 Hz. A 50 Hz digital notch filtering performed by the recording software was also used. R-peak detection was carried out automatically on the bipolar ECG-channel, then the RR-interval time-series extracted for each subject using the Kubios HRV software [Travainen et al. 2014]. Next, the Irregular-Resampling Auto-Spectral Analysis (IRASA) method [Wen et al. 2015] was applied to the entire time-series in the frequency range of 0.001–0.5 Hz, in order to separate the fractal and oscillatory PSD components, using a moving window of 4096 seconds.

Sleep structure indicators were available from the original study that included total sleep time, sleep efficiency, wake after sleep onset, sleep latency and relative durations of each sleep stage (W, N1, N2, N3, R).

### Spectral parametrization

As the fractal component exhibits a broken power-law relationship between the spectral power density and frequency, a piecewise linear function was fitted in the log-log domain, allowing for a custom breaking point, and two slope and intercept values in the lower and higher frequency domains.

The oscillatory component was defined as the log-log domain difference of the total PSD and the fractal component with gaussian smoothing applied (the standard deviation of the kernel was 20 samples). The peak frequency and prominence of the dominant peak was extracted by identifying local maxima from the LF (0.04–0.15 Hz) and HF (0.15–0.4 Hz) band, using the *find_peaks* function from the scipy.signal python library, see Figure 1.

**Figure 1.**
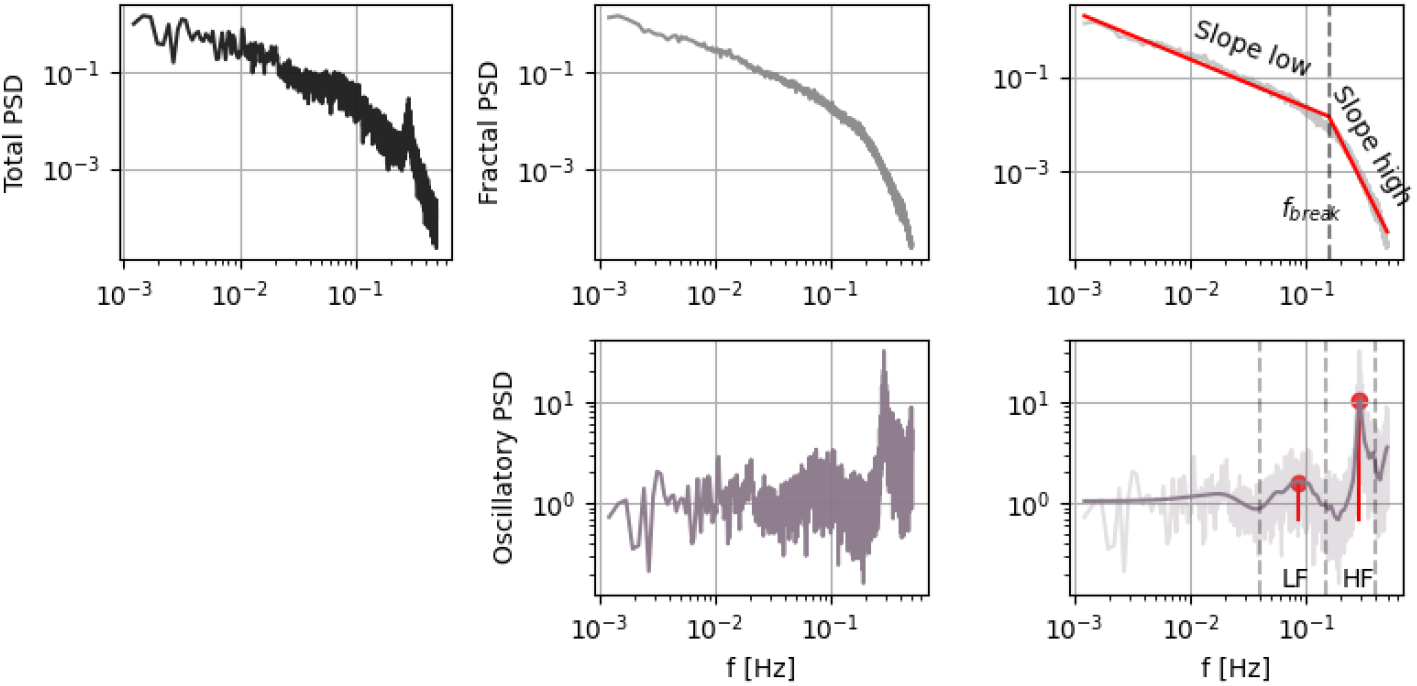
Outline of the spectral parametrization. The original PSD (top left) is split into the fractal and oscillatory components (middle column), then the fractal component is fitted with a piecewise linear function, allowing for a custom breaking point, that separates the linear fits in the low and high frequency domains (top right). Peak detection is applied in the LF and HF bands to the oscillatory component after gaussian smoothing, the frequency and prominence of the dominant peak is extracted from both bands (bottom right).

**Figure 2.**
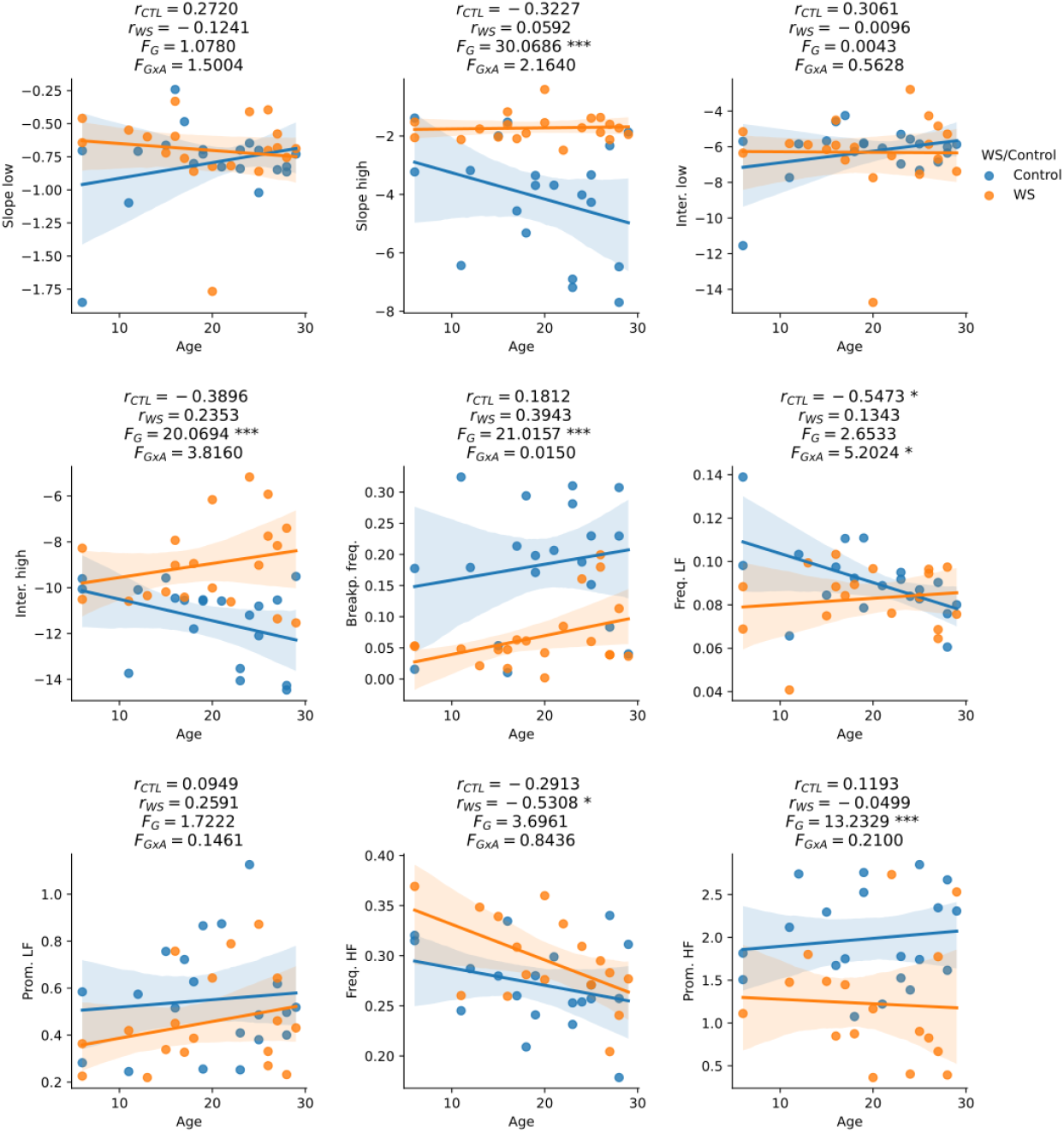
Effects of WS/TD group and age on the spectral parameters. Within group Pearson correlations with age are displayed above each plot for controls (r_CTL_) and WS (r_WS_) separately, as well as a general linear model results with main group effect statistic (F_G_) and the statistical power of the group*age interaction (F_GxA_). Significance marked as: *p<0.05, **p<0.01, ***p<0.001. (N=40 subjects, 20 WS)

### Statistics

A general linear model was assessed for each spectral parameter as the dependent variable including possible main effects of group (TD/WS), age and group*age interaction. Correlations between the spectral parameters and age were also calculated for each group separately.

As we revealed significant correlations between the parameters describing the fractal component, principal component analysis (PCA) was carried out to reduce the dimensionality of the data. Furthermore, the number of analysed sleep structure indicators was relatively large (nine in total), some of which were (by definition) correlated, thus PCA was applied in their case as well. As the dimensionality reduction proved to be meaningful both for the spectral and sleep structure parameters, associations between the two sets of principal components were further examined.

## Results

The broken power-law fitting was successful, mean goodness of fit was <R^2^>=0.990 (range=0.962-0.997, SD=0.007). Peaks were detected in all control subjects in both bands, whereas in the WS group 18 and 17 peaks were found in the LF and the HF band, respectively (out of the total 20 spectra, see Table 1. for more descriptive statistics).

**Table 1.** Group-wise descriptives of the broken power-law model parameters describing the fractal component (top) and the extracted peak parameters from the oscillatory components in the LF and HF bands (bottom).

|  | slope1 |  | slope2 |  | inter1 |  | inter2 |  | fbreak |  |
| --- | --- | --- | --- | --- | --- | --- | --- | --- | --- | --- |
|  | Control | WS | Control | WS | Control | WS | Control | WS | Control | WS |
| Valid | 19 | 20 | 19 | 20 | 19 | 20 | 19 | 20 | 19 | 20 |
| Missing | 0 | 0 | 0 | 0 | 0 | 0 | 0 | 0 | 0 | 0 |
| Mean | -0.804 | -0.700 | -4.168 | -1.722 | -6.298 | -6.309 | -11.473 | -8.968 | 0.183 | 0.068 |
| Std. Deviation | 0.312 | 0.296 | 1.999 | 0.443 | 1.521 | 2.309 | 1.697 | 1.839 | 0.102 | 0.054 |
| Minimum | -1.850 | -1.767 | -7.697 | -2.485 | -11.547 | -14.733 | -14.463 | -11.536 | 0.010 | 0.002 |
| Maximum | -0.242 | -0.331 | -1.385 | -0.406 | -4.247 | -2.784 | -9.511 | -5.174 | 0.324 | 0.199 |

|  | fpeak_lf |  | prom_lf |  | fpeak_hf |  | prom_hf |  |
| --- | --- | --- | --- | --- | --- | --- | --- | --- |
|  | Control | WS | Control | WS | Control | WS | Control | WS |
| Valid | 19 | 18 | 19 | 18 | 19 | 17 | 19 | 17 |
| Missing | 0 | 2 | 0 | 2 | 0 | 3 | 0 | 3 |
| Mean | 0.090 | 0.083 | 0.548 | 0.453 | 0.269 | 0.295 | 1.994 | 1.224 |
| Std. Deviation | 0.018 | 0.016 | 0.240 | 0.203 | 0.041 | 0.044 | 0.559 | 0.696 |
| Minimum | 0.061 | 0.041 | 0.245 | 0.219 | 0.178 | 0.204 | 1.075 | 0.363 |
| Maximum | 0.139 | 0.103 | 1.126 | 0.872 | 0.340 | 0.369 | 2.850 | 2.734 |

### Effects of WS group, age and interactions

Strong group effects were found in some of the fractal parameters, namely the high domain slopes were flatter (F_G_=30.07, p<0.001), high domain intercepts increased (F_G_=20.07, p<0.001), and the breaking point frequencies were decreased significantly (F_G_=21.02, p<0.001) in WS subjects compared to controls. An attenuation of HF peak prominences in WS (F_G_=13.23, p<0.001) was also observed.

A moderate slowing of LF peak frequency with aging (r=-0.55, p<0.05) was found in the control population, which however was not present in the WS group (r=0.13, NS), when assessing the correlations within the group, this difference was also confirmed by the group*age interaction in the general linear model (F_G*A_=5.20, p<0.05).

### Fractal parameters PCA

A high degree of correlation was found between specific pairs of fractal parameters (Table 2), which indicates that the variance in the data could be described by fewer parameters. In order to reduce the dimensionality of the fractal parameter space, PCA was applied, which resulted in two principal components. The first principal component was a linear combination of the high domain slope, high domain intercept and breaking point frequency, while the low domain slope and intercept contributed to the second principal component. Assessing the principal components as dependent variables in a general linear model as before, the main effect of WS/TD group was identified in the first component (F_G_=28.49, p<0.001), suggesting that WS-specific alterations in the RR-interval spectra are characterized by a joint increase in the high domain slope and intercept, along with a decrease in the breaking point frequency, as such, we will refer to this principal component as WS-related component, PC_WS_, see Figure 3. The second component was a linear combination of the low domain slope and intercept, which didn’t show effects of WS or age.

**Table 2.** Correlation matrix of all fractal parameter pairs with observations from all 40 subjects.

| Variable |  | slope1 | slope2 | inter1 | inter2 | fbreak |
| --- | --- | --- | --- | --- | --- | --- |
| 1. slope1 | Pearson's r | — |  |  |  |  |
|  | p-value | — |  |  |  |  |
| 2. slope2 | Pearson's r | 0.059 | — |  |  |  |
|  | p-value | 0.722 | — |  |  |  |
| 3. inter1 | Pearson's r | 0.914 | -0.114 | — |  |  |
|  | p-value | < .001 | 0.490 | — |  |  |
| 4. inter2 | Pearson's r | 0.104 | 0.825 | 0.043 | — |  |
|  | p-value | 0.527 | < .001 | 0.796 | — |  |
| 5. fbreak | Pearson's r | -0.011 | -0.883 | 0.200 | -0.538 | — |
|  | p-value | 0.947 | < .001 | 0.223 | < .001 | — |

**Figure 3.**
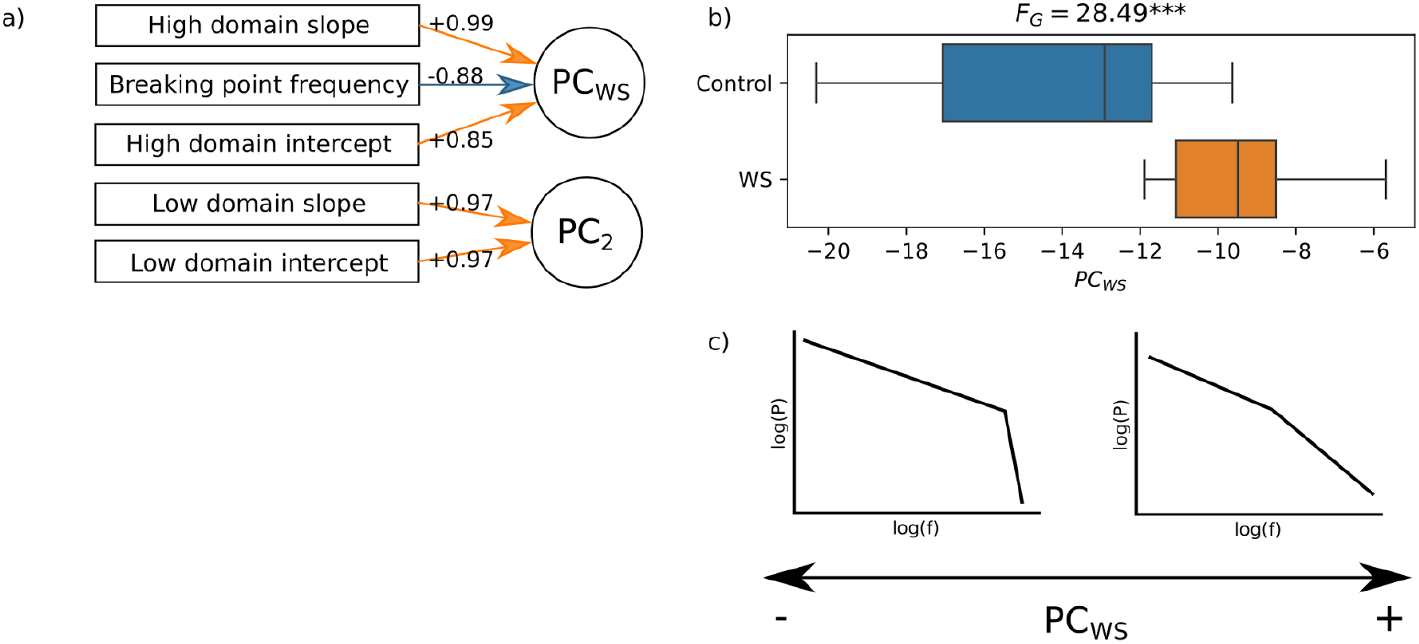
Principal component analysis in the fractal parameters. Due to the high correlation of the original parameters, the main WS vs TD group differences in the fractal spectra can be described by a single component PC_WS_. a) Contributions of the fractal parameters to the two principal components. b) The first component shows a significant effect of WS. c) Illustration of characteristic changes in the power spectrum explained by the PC_WS_ component.

### Sleep structure PCA

Many pairs of the nine sleep structure indicators were also significantly correlated (15 out of 36 total pairs), which were reduced by PCA to two components (Chi^2^=1007.89, p<0.001). The first component had positive contributions of sleep efficiency, sleep duration and relative REM duration and negative contributions of relative wake duration, wake after sleep onset and sleep latency, so it can be termed as a general sleep-quality component PC_SQ_. The second component is positively associated with relative SWS duration, and negatively with relative NREM2 and NREM1 durations, and thus an indicator of deep versus light sleep duration ratio, referred to as sleep depth component PC_SD_, see Figure 4.

**Figure 4.**
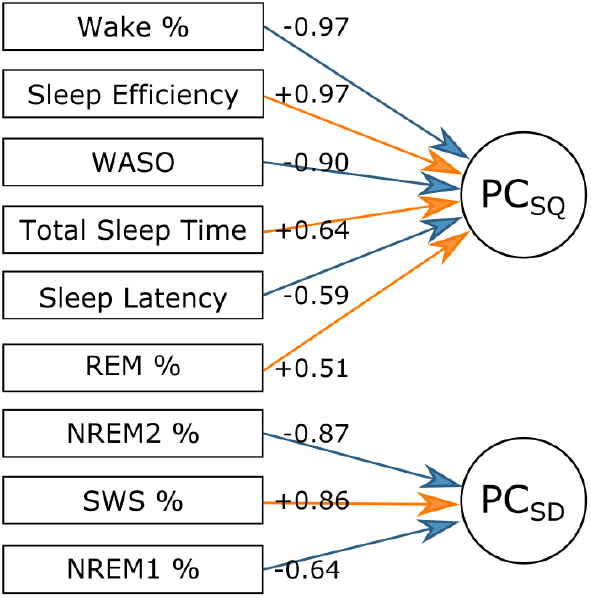
Principal component analysis of sleep structure indicators. Two main components were identified. Higher values of the first component could be associated with better sleep quality PC_SQ_, while the second increased with deep to light sleep duration ratio PC_SD_.

After the reduction of the parameters, the correlations of the PC_WS_ component were assessed with the PC_SQ_ and PC_SD_ components. Higher Williams component scores were associated with significantly lower sleep-quality PC_SQ_ components (r=-0.355, p<0.05) and increased PC_SD_ sleep depth component values (r=0.440, p<0.01), see Figure 5.

**Figure 5.**
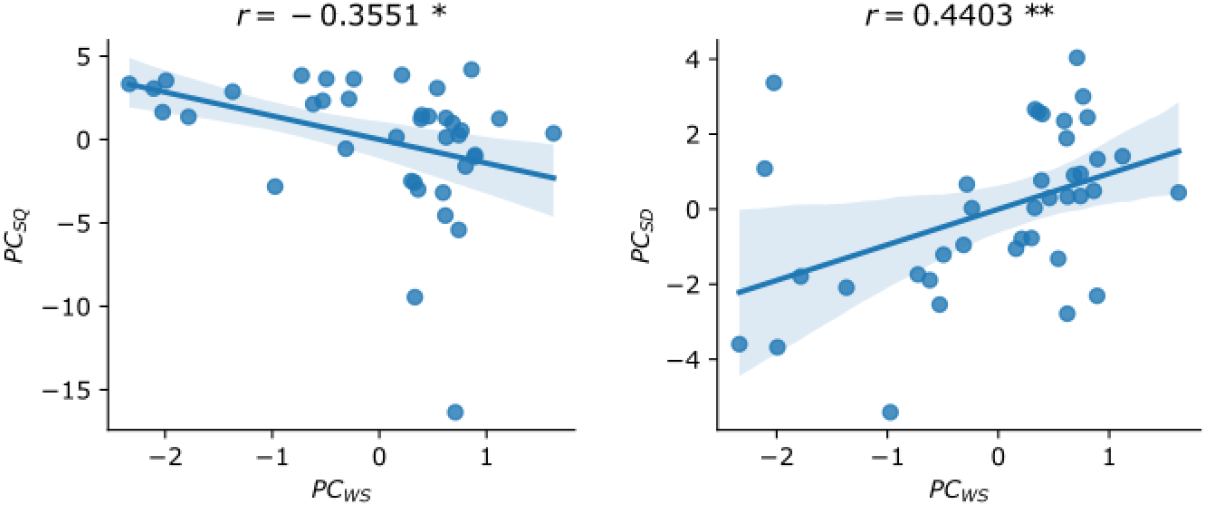
The PC_WS_ component is correlated with the principal components of the sleep indicators, increased PC_WS_ scores imply worse sleep quality and a shift toward deeper sleep. *p<0.05, **p<0.01, N=40 subjects.

## Discussion

One of our main findings was that the parameters of the broken power-law model describing the fractal component of the HRV spectrum were altered in WS. More specifically, the breaking point frequency was reduced (shifted toward the lower frequencies) and an increase in the slope (flattening of the high frequency spectra) and intercept above the breaking point were revealed. An apparent contradiction between our findings and the general power reduction reported in a previous study [Levin et al. 2023] might arise, however, we note that there were no significant group differences in the fractal parameters below the breaking point, and slopes above it were steeper in general in both groups (see Table 1.), thus lower breaking point frequencies imply less total power. At the same time, the mean breaking frequency in WS was at 0.06 Hz, well below the upper boundary of the LF band (0.15 Hz), meaning that the changes in the fractal component can explain power reduction both in the LF and HF bands. In addition, the attenuation of HF peak prominence can also be responsible for the increased LF/HF ratio reported in the same study.

The phenomenon of accelerated aging [Bódizs et al. 2012, 2014] was also reflected in the LF peak frequency, as it showed a significant deceleration with age in the control group, but generally lower values regardless of age in WS, also confirmed by the interaction term in the GLM.

We propose that the flattening of the high frequency spectral slope could reflect the erratic, beat-to-beat type fragmentation of heart rate in WS subjects reported recently (Cathey et al, 2024). Indirect evidence supporting this assumption can be based on the fact that heart rate fragmentation is quantified on the basis of the relative density of inflections in the RR-series. Frequent inflections indicate frequent short scale changes in the time series, which are known to be reflected in a flatter spectral slope. More specifically, the nominal value of the group mean of high frequency spectral slope in WS subjects (−1.722) falls in the so-called antipersistent range (>−2), whereas in the TD participants the same value (−4.168) is definitely below the threshold of −2, indicating a persistent time series. The former reflects rapid shifts in the heart rate increments (decreases tend to be followed by increases and vice versa), whereas the persistent spectra implies a positive correlation in the TD subjects (increases tend to be followed by further increases and vice versa) [Carpena et al., 2022]. Present results suggest that bifractal spectral examination of heart rate is potentially indicative of the erratic cardiovascular dynamics in WS, which were formerly proposed to reflect the effects of non-autonomic cardiovascular conditions (Cathey et al., 2024).

In addition to the alterations of fractal HRV spectra outlined above, the decrease of HF peak prominence in WS subjects was also revealed. Decreased HF peak prominence indicates a reduction in respiratory sinus arrhythmia and a related reduction in vagal modulation of heart rate of WS subjects during sleep. Together these findings indicate that the heart rate regulation during human sleep is a result of a complex combination of bifractal and oscillatory processes. The combined alteration of the respective measures in WS suggest that autonomic and cardiovascular modulators interact in shaping the heart rate dynamics of WS subjects during sleep.

Moreover, the present results further strengthen the pluripotent clinical value of polysomnographic studies. The latter involve an ECG derivation, thus are ideally suited for revealing the short- and long-term dynamics of resting heart rate during sleep in various clinical conditions and research settings [Stein and Pu, 2012].

As the principal component analysis successfully reduced the correlated fractal parameters to only two components, a general feature could be identified in the HRV spectra, typical of individuals with WS. Furthermore, this WS component was negatively correlated with sleep quality indicators and positively associated with higher deep to light sleep ratio, indicating that it is not only a feature of cardiac rhythm alteration, but of general deregulation in the autonomic nervous system, which couples sleep homeostasis with cardiovascular control, thus a potential biomarker for WS severity.

## Funding

Supported by the University Research Scholarship (2024-2.1.1-EKÖP-2024-0004), as well as the Thematic Excellence (TKP2021-EGA-25 and TKP2021-NKTA-47) Programmes of the Ministry for Culture and Innovation from the source of the National Research, Development and Innovation Fund, Hungary. It was made in the framework of the PPKE-BTK-KUT-23-1 project, with the support and funding provided by the Faculty of Humanities and Social Sciences of Pázmány Péter Catholic University.

## Notes

### Competing Interest Statement

The authors have declared no competing interest.

